## Supplementary Information for "Basolateral amygdala oscillations enable fear learning in a biophysical model"

**VIP and SOM interneurons fire at gamma nested low theta and high theta, respectively.**

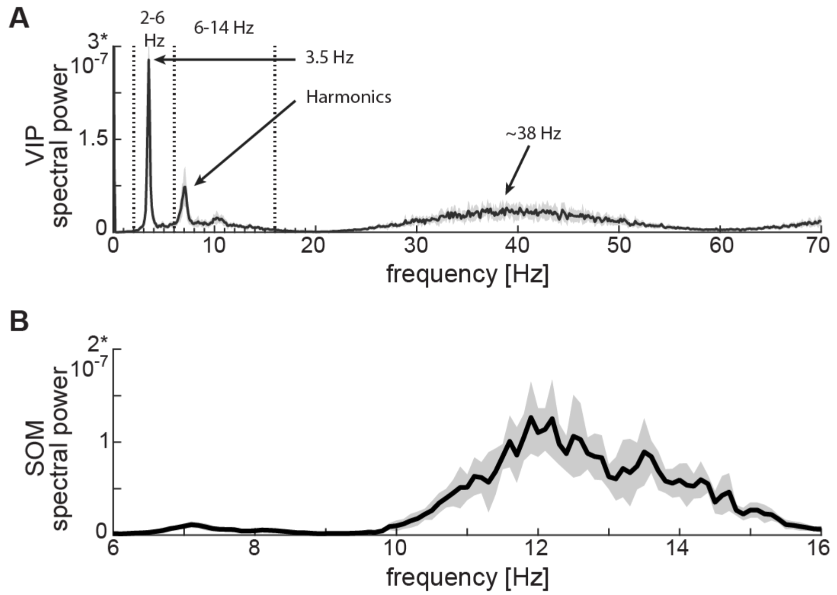


**Figure S1**. **Power spectra of neuronal spiking activity at baseline. A**: Power spectrum of VIP cell showing a first peak at low theta (~3.5 Hz), plus harmonics, and a second peak at gamma (~38 Hz). **B**: SOM cell power spectrum showing a peak at high theta (~12 Hz). Both panels show the mean (black curves) and standard deviation (black shaded areas) of the power spectra across 10 network realizations.

**ECS and F activity patterns determine overall potentiation or depression.** The STDP model used here considers the whole history of ECS and F spikes by exploiting functions P, used for potentiation, and M, used for depression, as defined in (Song et al., 2000). P evolution in time is determined by ECS (presynaptic neuron) spiking activity, while M is shaped by F (postsynaptic neuron) spiking activity. As detailed in the section "Synaptic plasticity" in the Materials and Methods, these two functions change at the time a neuron spikes and then relax exponentially towards the equilibrium (0) (see Fig. S2).

When both ECS and F are firing at gamma (as shown in Fig. S2B, right, in the first 100 ms), both P and M build up, but since M has a longer relaxation time it builds up more than P, such that after ~100 ms of gamma depression dominates. Note that M and P decay back to zero if there is a pause in spiking for a theta cycle (Fig. S2B, right, in between ~8150-8250 ms). The actual change of synaptic conductance is computed as follows. When F spikes, the instantaneous value of P determines the amount of potentiation the synapse from ECS to F undergoes. By contrast, when ECS spikes, the instantaneous value of M determines the amount of depression that weakens that synapse. Fig. S2 presents the P and M functions, along with the network dynamics and the evolution of the conductance from ECS to F, for the full network (Fig. 2), the only-PV network (Fig. 4A), and the PV/VIP network (Fig. 4B) in the presence of CS and US.

By using a depression-dominated rule (Fig. S2A), we show in the main text that the ECS to F conductance overall potentiates in presence of the full network after the US onset (Fig. 2). Fig. S2B shows that overall potentiation wins over depression because: (i) during the long disinhibition windows at low theta provided by VIP, both ECS and F are active and ECS fires most of the time slightly before F; despite M being slightly stronger than P, the pre-post fine timing ensures potentiation; (ii) M relaxes towards the equilibrium during the silent ECS-F phase of the low theta rhythm, and thus the potentiation acquired during the active ECS and F phase builds up over time.

In the PV-only network (Fig. S2C), with PV in a low excitation regime such that it can entrain with F in a coordinated gamma rhythm (PING), we show in the main text that there is overall depression in the conductance from ECS to F (Fig. 4A) because PING is not periodically interrupted. Indeed, PING without interruptions makes the M function saturate. By contrast, the P function shows only a few jumps and long periods at the equilibrium due to a few ECS spikes. Thus, a weak potentiation happens at each F spike, but it is counterbalanced by the strong depression at each (despite few) ECS spike, leading to overall depression.

In the network with PV and VIP (Fig. S2D), we find that VIP cell reduces the excitation of the PV cell (which is set at high excitation, unless otherwise specified) during its active low-theta phase, enabling the PV cell to participate in PING. Secondly, it provides periodic interruptions in PING (Fig. 4B). Fig. S2D shows that potentiation in the conductance from ECS to F arises during the active VIP phases because of the ECS-F fine timing. However, during the silent VIP phases, ECS is still active due to the weaker inhibition from PV and the lack of inhibition from SOM. At each ECS spike, a non-negligible depression happens despite the M function relaxing. In conclusion, the depression brought about by ECS spikes during the silent VIP phases counteracts the potentiation acquired during the active VIP phases such that, overall, no significant potentiation takes place.

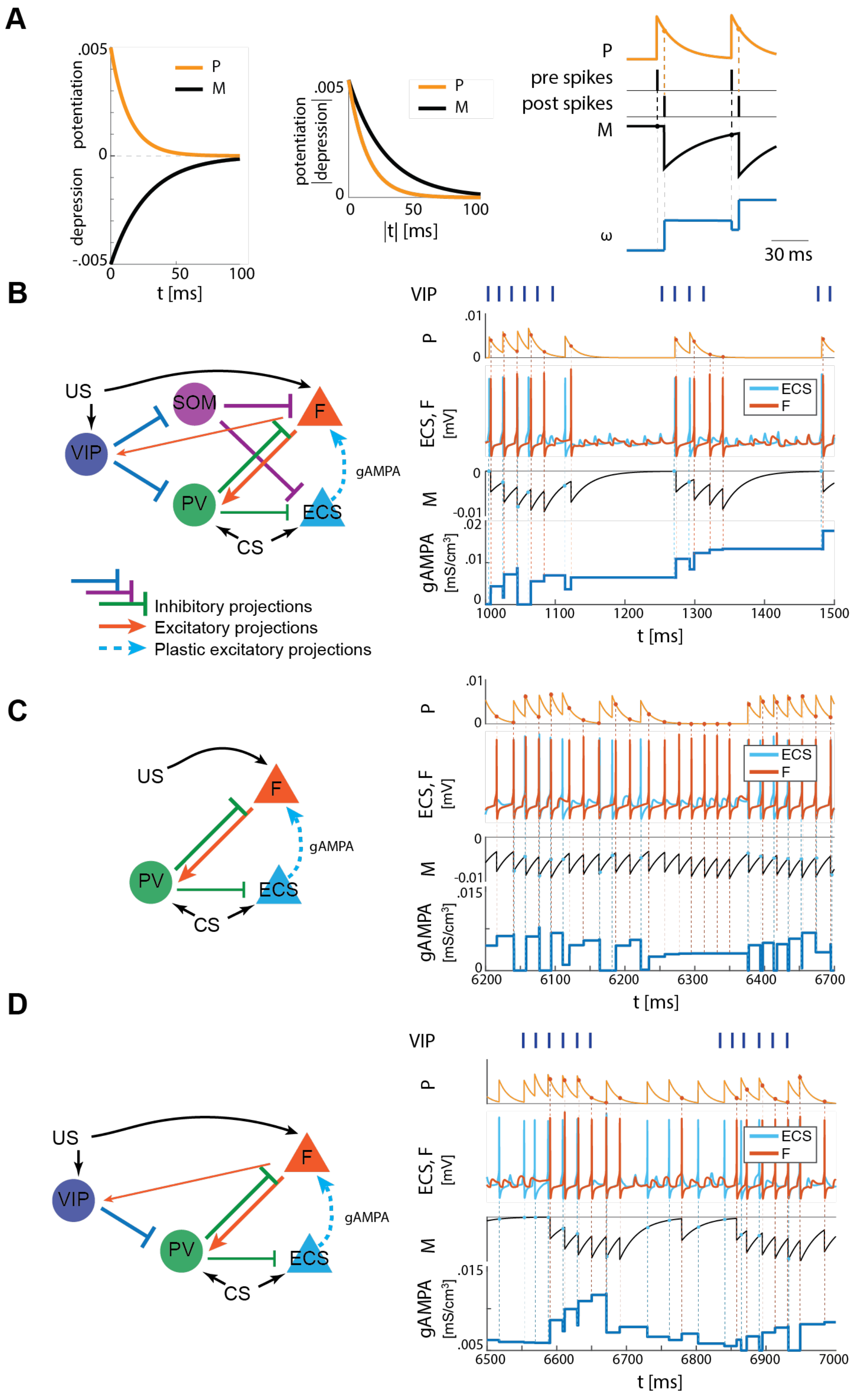


**Figure S2**. **Plasticity rule and detailed representation of how plasticity works in the three network configurations shown in Fig. 2B and Fig. 4A,B. A**: Left, STDP potentiation (P) and depression (M) functions. The area underlying depression is larger than the one of potentiation, thus providing a depression-dominated rule. The potentiation and depression curves are used to compute the instantaneous change of the synaptic conductance as a function of the spike time difference between pre and postsynaptic neurons, as detailed in the section "Synaptic plasticity" in Materials and Methods, and in the Supplementary Information. Right, a representative spike pattern of pre and postsynaptic neurons alongside the resulting P and M functions and the evolution of the pre-post synaptic conductance. **B**: Left, full network. Right, ECS (pre) and F (post) spiking activities over 500 ms (extracted from Fig. 2) with their respective M and P functions, which determine how the AMPA conductance from ECS to F evolves in time. **C**: Left, only-PV network at a low excitation level. Right, ECS and F dynamics as in 500 ms extracted from Fig. 4A with P, M, and AMPA conductance unfolded over time. **D**: Left, network with VIP and PV. Right, ECS and F dynamics as in 500 ms extracted from Fig. 4B, alongside P, M, and AMPA conductance over time.

**An alternative network configuration characterized by US input to PV, instead of CS, also learns the association between CS and fear.** We constrained the BLA network in Fig. 2 with CS input to the PV interneuron, as reported in (Krabbe et al., 2018). However, (Krabbe et al., 2019) notes that a class of PV interneurons may be responding to US rather than CS. Fig. S3 presents the results obtained with this variation in the model (see Fig. 3 A,B for comparison) and shows that all the network realizations learn the association between CS and fear. In the model, the PING rhythm between PV and F is the crucial component for establishing fine timing between ECS and F, which is necessary for learning. Having PV responding to the same input as F, i.e., US, facilitates their entrainment in PING and, thus, successful fear learning.

We model the VIP interneuron as affected by US; in addition, (Krabbe et al., 2019) reports that a substantial proportion of them is mildly activated by CS. Replacing the US by CS does not change the input to VIP cells, which is modeled by the same constant applied current. Thus, the VIP CS-induced activity is a bursting activity at low theta, similar to the one elicited by US in Fig. 2.

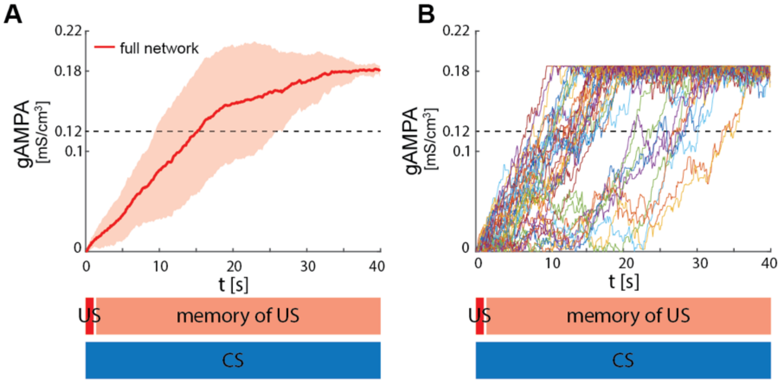


**Figure S3. ECS to F conductance across network realizations in BLA characterized by US, instead of CS, input to PV interneuron.** **A**: Mean (color-coded curves) and standard deviation (color-coded shaded areas) of the AMPA conductance (gAMPA) from ECS to F across 40 network realizations over 40 seconds. **B**: Evolution in time of the AMPA conductance for the 40 full network realizations in A.

**Classical Hebbian plasticity rule, unlike the depression-dominated one, shows potentiation even with no strict pre and postsynaptic spike timing.**

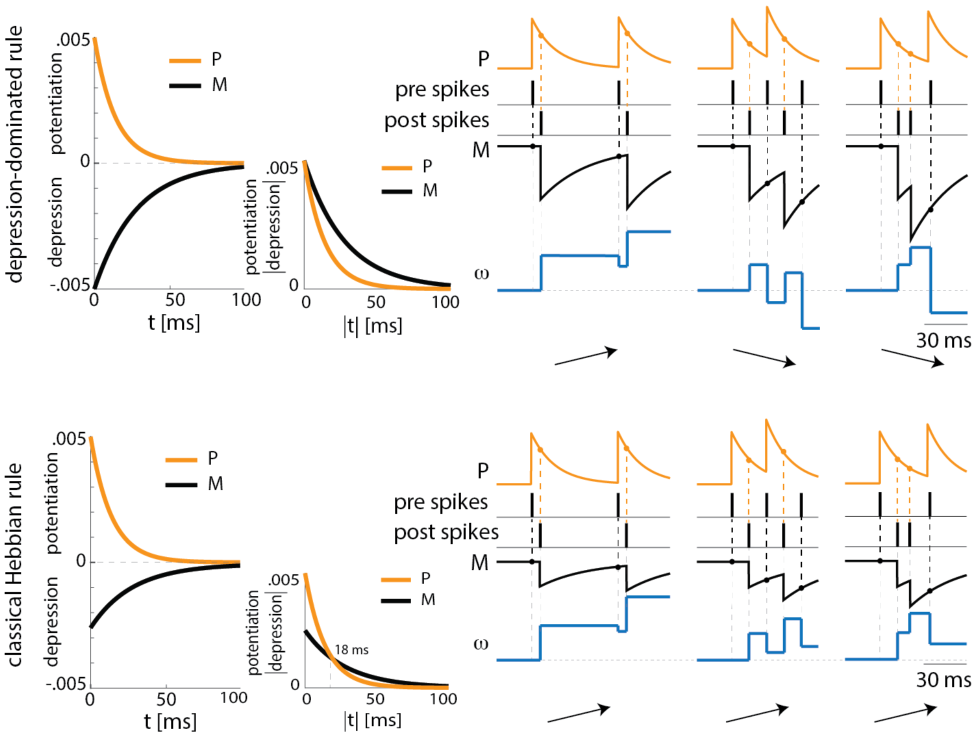


**Figure S4**. **Depression-dominated and classical Hebbian plasticity rules may provide opposite potentiation/depression profiles.** Top, depression-dominated rule (as in Fig. S2A) with three examples of pre and postsynaptic spike patterns. Only the first one, which shows a consistent pre-post timing, shows overall potentiation. The remaining two spike patterns lead to depression. Bottom, classical Hebbian rule characterized by a smaller maximal amplitude for depression than potentiation. In agreement with the depression-dominated rule, the classical rule shows potentiation in the case of the pre and postsynaptic neurons showing correct pre-post spike timing. However, the classical rule shows potentiation also in the remaining two examples where there is no-consistent pre-post timing, given that the pre and postsynaptic neurons fire at a frequency higher than 55 Hz.

**A higher low theta power increase emerges in LFP approximated with the sum of the absolute values of the currents compared to their linear sum.** Given that our BLA network comprises a few neurons described as single-compartment cells with no spatial extension and location, the LFP cannot be computed directly from our model's read-outs. In the main text, we choose as an LFP proxy the linear sum of the AMPA, GABA, and P-/H-/D-currents. We note that if the LFP is modeled as the sum of the absolute value of the currents, as suggested by (Mazzoni et al., 2008, 2015), an even higher low theta power increase arises after fear conditioning compared to the linear sum. Differences in the power spectra also arise if other LFP proxies (e.g., only AMPA currents, only GABA currents) are considered. A principled description of an LFP proxy would require modeling the three-dimensional BLA anatomy, including that of the interneurons VIP and SOM; this is outside the scope of the current paper. (See (Feng et al., 2019) for a related project in the BLA.)

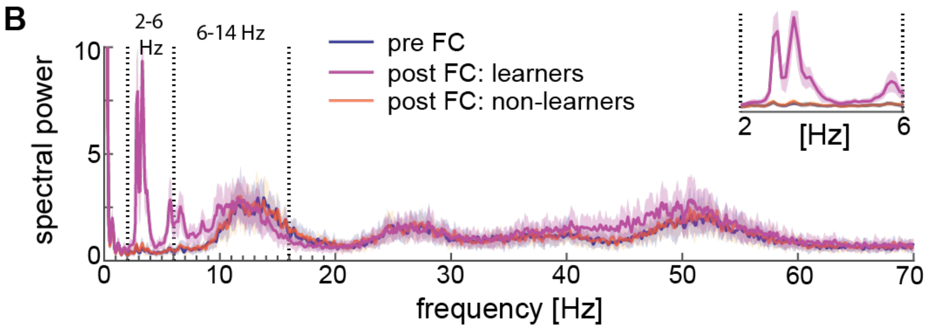


**Figure S5**. **Spectral properties pre versus post fear conditioning of LFP approximated with the sum of the absolute values of AMPA, GABA, D-, NaP-, and H-currents.** Power spectra before fear conditioning (blue) and after successful (purple) and non-successful fear conditioning (orange); top, right: inset between 2 and 6 Hz. Blue and orange curves closely overlap.
